# Fundamental resource specialization of herbivorous butterflies decreases toward lower latitudes

**DOI:** 10.1101/2020.01.09.899922

**Authors:** Ryosuke Nakadai, Tommi Nyman, Koya Hashimoto, Takaya Iwasaki, Anu Valtonen

## Abstract

**Aim:** It is generally assumed that the degree of resource specialization in herbivorous insects increases towards lower latitudes. However, latitudinal patterns in herbivore diet breadth at large spatial scales remain poorly understood. In this work, we investigated drivers of latitudinal variation in lepidopteran “fundamental” resource specialization, which we defined as the host breadth when not limited by interspecific interactions at the same trophic level.

**Location:** The Japanese archipelago (22°N–45°N), including hemiboreal, temperate, and subtropical zones.

**Taxon:** Herbivorous butterflies.

**Methods:** Species-specific fundamental host breadth was calculated based on pooled geographical occurrence and host-use records. We investigated the latitudinal pattern and significant drivers of the degree of specialization in regional species pools at a 10-km grid level. As potential drivers, we focused on geography, current climate, and diversity and body size of butterflies. Through Bayesian structural equation modeling, we investigated the complicated relationships between these variables and community-level resource specialization represented by three different indices of host breadth.

**Results:** We found that fundamental resource specialization of butterfly communities increases toward higher latitudes. This pattern is contrary to the presumed general trend found in studies based on realized resource specialization within local communities. We found that the observed pattern is driven mainly by factors related to climate, butterfly diversity, and body size in each community. Above all, annual mean temperature most strongly drove community-level fundamental host breadth of butterflies.

**Main conclusions:** Our findings suggest that fundamental resource specialization may show different latitudinal patterns from the conventional prediction based on knowledge of realized resource specialization. Our results emphasize the importance of the current climate as a major factor regulating butterfly morphology and fundamental host breadth, regardless of whether the impact is direct or indirect.

## Introduction

Understanding the processes structuring communities across space and time is a central subject of ecological research (Cavender-Bares et al., 2009). Resource specialization is a major factor shaping interspecific interactions and, therefore, community composition (Nylin et al., 2017; Volf et al., 2017; Dyer & Forister, 2018). The drivers of variation in the degree of specialization have been intensively investigated in the context of community ecology (Novotny & Basset, 2005; Forister et al., 2015). Such studies have generally reported that species tend to be more specialized in the tropics than in the temperate zone (MacArthur’s latitude–niche breadth hypothesis; MacArthur, 1972; Vázquez & Stevens, 2004; Dyer et al., 2007; reviewed by Dyer & Forister 2018). Using a global dataset, Forister et al. (2015) addressed variation in resource specialization of herbivorous insects along a geographical gradient, and showed that the mean and variance of diet breadth in local communities decrease toward the equator.

Resource specialization is driven not only by geographical and climatic gradients, but also by many other factors. Specifically, the higher number of species in the tropics (Loder *et al*., 1998; Dyer & Forister, 2018) may intensify interspecific competition, thereby favoring “species packing” (Dobzhansky, 1950; MacArthur, 1972) via specialization (Coley & Kursar, 2014). The importance of body size and physiological constraints imposed by climatic conditions has long been recognized (Bergmann’s rule; Bergmann, 1847). For example, average consumer body size and its variation increase toward lower latitudes (Loder et al., 1998), although body size is generally positively correlated with host breadth (Lindström et al., 1994; Pöyry et al., 2017). Identification of the drivers of specialization is further complicated by the fact that numerous potentially important factors change in parallel along latitudinal and climatic gradients (Loder *et al*., 1998).

On the other hand, some studies have found no differences in the degree of specialization between temperate and tropical regions (e.g., Novotny et al., 2006), and the hypothesis remains highly controversial (Moles & Ollerton, 2016). To date, most studies on variation in resource specialization have relied on data on local communities. However, resource use within local communities can be shaped by direct or indirect interspecific interactions (Connell, 1980), and local processes of community assembly can produce “realized” resource specialization, which may mask underlying fundamental resource specialization. Shifting the focus to fundamental resource specialization can therefore facilitate understanding processes driving variation in the degree of resource specialization without considering the complicated effects of local interspecific interactions. In this work, we investigated the processes driving spatial variation in “fundamental” resource specialization, which we define as host breadth when not limited by interspecific interactions at the same trophic level. More specifically, we aimed to reveal the drivers of geographical variation in fundamental resource specialization using macroecological approaches.

We focus on angiosperm-feeding butterflies across the Japanese archipelago to reveal the drivers of spatial variation in resource specialization. We calculated species-specific fundamental host breadths based on pooled geographical occurrence data and host-use records. The latitudinal range of the archipelago (22°N–45°N) includes hemiboreal, temperate, and subtropical zones, which makes it a useful system for assessing spatially continuous changes in resource specialization, rather than a comparison between disconnected areas. In fact, most previous studies have simply compared resource specialization between tropical and temperate regions, which Dyer and Forister (2018) highlighted as a major methodological problem. We focused on the latitudinal pattern and aimed to reveal significant drivers of the degree of specialization in regional species pools. As potential drivers, we focused on geography, current climate, diversity and body size of butterflies, and their changes along geographical gradients (Table 1). We identified drivers of resource specialization using Bayesian structural equation modeling (Bayesian SEM), which has received increasing attention in ecology and evolution (e.g., Arhonditsis et al., 2007; Read et al., 2018; reviewed in Grace et al., 2012). Specifically, we investigated the complicated relationships between the explanatory variables and resource specialization using a comprehensive model fitted with a single index reflecting the distribution of host breadth, and the mean and divergence of phylogenetic host breadth within each community.

**Table 1.**
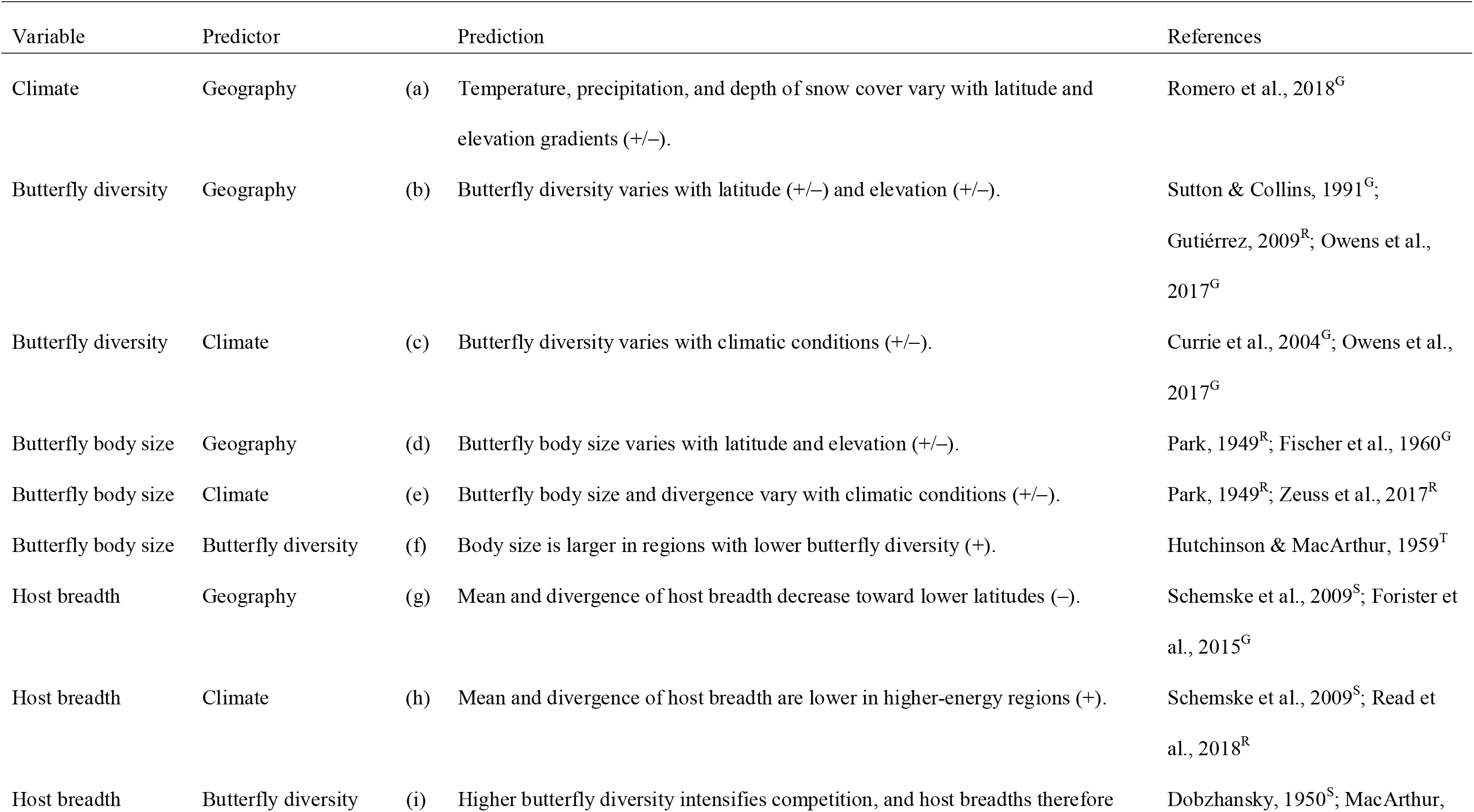
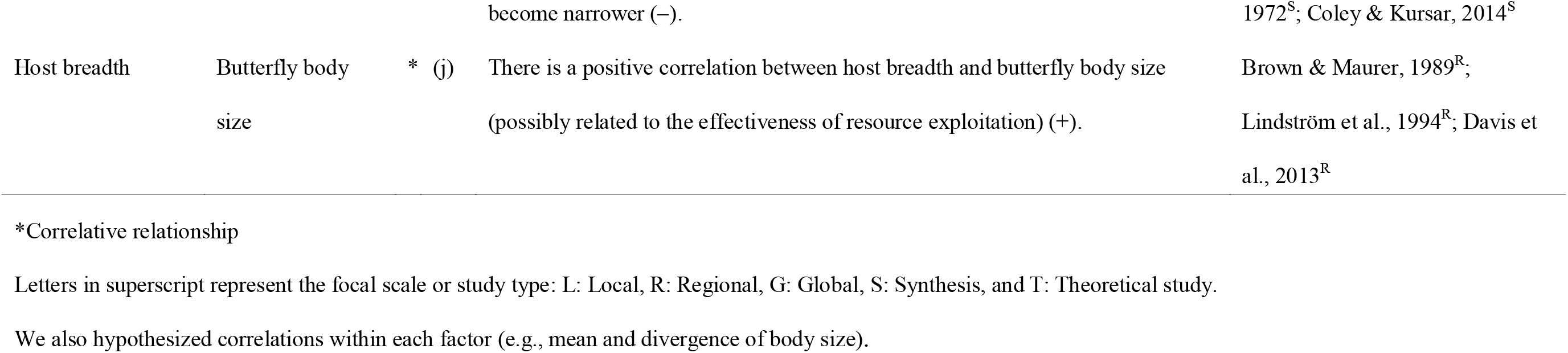
Hypothetical trends observed in previous studies and considered in this study.

## Materials and Methods

### Study area

The Japanese archipelago is a long chain of continental islands located off the eastern coast of Asia, and is recognized as a hotspot of biodiversity (Mittermeier et al., 2011). The mean temperature of the coldest month ranges from −18.4°C to 22.3°C, while that of the warmest month ranges from 6.2°C to 29.2°C, and annual precipitation is 700.4–4477.2 mm (Fig. S1a, b: Japan Meteorological Agency, 1981–2010; http://www.jma.go.jp/jma/indexe.html). Northern Japan, particularly Hokkaido, receives heavy snowfall during the winter. By contrast, the Ryukyu Islands, a subtropical region in the southern part of the archipelago, are characterized by warm temperatures year-round, with little seasonal variation. The western (Sea of Japan) side of the archipelago typically receives more snow than the eastern (Pacific Ocean) side (Fig. S1c). The Japanese archipelago belongs to a transitional region between the Paleotropical or Indo-Pacific and Holarctic regions (Cox, 2001), and harbors relict plants that were distributed throughout a large portion of the Northern Hemisphere during much of the Tertiary (Milne & Abbott, 2002). More than 280 species of butterflies in five families are found in the region (Shiro□zu, 2006). With the exception of three lycaenid species, larvae of Japanese butterfly species feed exclusively on plants (Honda, 2005). Their host plants are diverse and include both dicot and monocot angiosperms, as well as gymnosperms (Saito et al., 2016).

### Evaluation of butterfly host breadth and body size

Our analyses were based on butterfly census data, including records of 273 species or subspecies of butterflies throughout the entire Japanese archipelago, available from the website of the Biodiversity Center of Japan, Ministry of the Environment (http://www.biodic.go.jp/index_e.html), in grid cells of 5 min latitude and 7.5 min longitude (about 10 km × 10 km; the Japanese Standard Second Mesh). We used the results of the fourth and fifth censuses (1988–1991 and 1997–1998, respectively) of the National Survey of the Natural Environment in Japan (http://www.biodic.go.jp/kiso/15/do_kiso4.html). This database includes records from 273 species or subspecies of butterflies in grid cells throughout the Japanese archipelago. We aggregated data at the species level and converted all records to presence versus absence (1/0) of each species in each grid cell. Following aggregation, 93,449 records, across all targeted species, were used in the study. For each species, records were used in combination with relevant climate data to predict potential distributions throughout Japan using Maxent ver. 3.3.3e (Phillips et al., 2006; analyzed in Nakadai et al., 2018). To compensate for incomplete distributional data, distribution probabilities were converted to potential distributions (0/1) using maximum training sensitivity plus specificity logistic thresholds. Due to difficulties in estimating the potential distributions of rare species using Maxent (Vaughan & Ormerod, 2005; Breiner et al., 2015), the presence records of 18 butterfly species with four or fewer records each were used directly in subsequent analyses.

Information on the host plants of each studied butterfly species was compiled for evaluation of diet breadth. The host plants of 278 butterfly species or subspecies (3,573 records in total, after excluding duplicates) were obtained from Saito et al. (2016). We used adult forewing length as a proxy for the body size of each butterfly species. The forewing lengths of 284 species were derived from Shiro□zu (2006; published in Nakadai et al., 2020). For a total of 247 butterfly species, we assembled data on predicted distribution, host plants and body size. However, nine species were excluded from the analyses for various reasons (see Table S1). After the non-target species (those with taxonomic changes or non-angiosperm hosts) were removed, the remaining 3,377 host plant records were converted into plant-genus resolution for further analyses.

To describe the overall degree to which species are specialized in their host use, we estimated the alpha parameter of the discrete truncated Pareto distribution fitted to numbers of host genera for all butterfly species occurring within each grid cell (Forister et al., 2015; Kozubowski et al., 2015). For highly nonsymmetrical distributions, the shape parameter (α) of the discrete truncated Pareto distribution is more informative than measures of central tendency such as the mean (Forister et al., 2015). Higher values of the Pareto alpha parameter indicate increased specialization in the community, i.e., a higher proportion of species with relatively narrow host breadth.

For each cell, we also calculated the grid-level community mean and divergence (based on Rao’s quadratic entropy; RaoQ) of both phylogenetic host breadth and butterfly body size using the ade4 package (Chessel et al., 2013) in R (R Development Core Team 2017). The community mean indicates the dominant trait value within an assemblage (Violle et al., 2007), while community divergence represents the degree of dissimilarity between all possible pairs of species within the assemblage (Mouchet et al., 2010). To describe the genus-level phylogenetic host breadth of butterfly species, we applied Faith’s phylogenetic diversity index (cf., Davis et al., 2013). We calculated this index with the R package picante (Kembel et al., 2010) based on a genus-level angiosperm phylogeny derived from Zanne et al. (2014) using Phylomatic (Webb & Donoghue, 2005). The methods for estimating phylogenetic host breadth and the distribution of host breadth followed those of Forister et al. (2015), which provides detailed descriptions and statistical background information.

To determine whether the observed mean and divergence of each trait in each grid were greater or less than the values obtained by drawing butterfly species at random from the regional species pool, we calculated the standardized effect size (SES) using two types of null models (i.e., null model 1 and 2). “Null model 1” was generated by sampling the grid cell-specific number of species, without replacement, from the pool of all 247 focal species. This null model standardizes the values of the indices by removing bias resulting from differences in the number of species within grid cells (Mason et al., 2007). “Null model 2” was generated by randomization of the grid cell–species matrix using an independent swap algorithm (Gotelli, 2000), thereby retaining the original number of species in each grid cell, as well as the frequency of occurrence of each species across all grid cells. The randomization procedures for both null models were repeated 1,000 times to generate simulated datasets, which were then used to compute cell-specific expected values for the mean and divergence of host breadth and body size. We calculated SES values for each grid cell as the observed value minus the mean of the null distribution, and the result was divided by the standard deviation of the null distribution. Therefore, a negative RaoQ value indicates that trait values within the cell are more similar than expected based on the null model (i.e., convergence within the community), whereas a positive value indicates that species in the assemblage are more dissimilar than expected (i.e., divergence within the community). For our main analyses, we used datasets based on the simpler null model 1, and then re-fitted the final model using null model 2 to explore the robustness of our results.

### Climatic variables

We used annual mean temperature and precipitation at the 1 km grid scale from Mesh Climate Data 2010 (Japan Meteorological Agency, http://nlftp.mlit.go.jp/ksj/gml/datalist/KsjTmplt-G02.html). We also included annual maximum depth of snow cover (Mesh Climate Data 2010), and elevation data from the National Land Numerical Information database (Geospatial Information Authority of Japan). All of these variables were downloaded at the 1 km grid scale. Finally, for all variables, we calculated average values at the 10 km grid scale using ArcGIS Pro ver. 2.3.3 software (Environmental Systems Resource Institute, Redlands, CA, USA).

### Driver analyses for host breadth and body size

To reveal the drivers of resource specialization, we compiled a Bayesian SEM in which the alpha parameter of the discrete truncated Pareto distribution and the two indices (i.e., mean and divergence) of phylogenetic host breadth were explained by butterfly species richness, two indices of body size (i.e., mean and divergence) and five environmental factors. The environmental factors included as explanatory variables were latitude and elevation as geographical factors, and annual precipitation, mean temperature and annual maximum depth of snow cover as climatic factors. Although climatic and geographical factors are correlated (Table S2), including both types of correlated variables may be informative. For example, a strong negative correlation exists between latitude and annual mean temperature, but other effects of latitude may include ecological factors that were not considered in this study, such as variation in day length and the length of the active season.

Values of *fit* were calculated by the average least-squares differences between the values of observed and predicted host breadth as 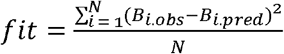, where *B*_*i*.*obs*_ and *B*_*i*.*pred*_ are the relative numbers of species with host breadth *i* and N is the maximum host breadth in a community. Specifically, *B*_*i*.*obs*_ and *B*_*i*.*pred*_ are calculated as the number of species with host breadth *i* divided by total species richness in a community. If the total values of observed and predicted species richness are almost the same, a *fit* ≥ 0.01 indicates that the deviation between the observed and predicted number of species of each host breadth is more than ten percent on average. Prior to analyses, two types of grid cells were excluded from the dataset following Ulrich et al. (2016): cells with fewer than 10 butterfly species and cells with *fit* ≥ 0.01. These cells (N = 378 and N = 695, respectively: N = 759 in total) were excluded because their values were considered less reliable or due to difficulty in correctly estimating the alpha parameter of the discrete truncated Pareto distribution. For the first criterion, we repeated the analyses with the minimum number of species set to 15 or 20 to validate the robustness of the main results.

We fitted the full Bayesian structural equation model in Mplus version 8.4 (Muthén & Muthén, 1998–2017). Starting parameters were selected using the maximum likelihood method (Muthén & Muthén, 1998–2017). We ran three Markov chain Monte Carlo analyses with 1,000,000 iterations from the posterior distribution, using the Metropolis–Hastings algorithm. Every 100th iteration was sampled from each chain, and the first half of the sample set was discarded as burn-in, which is the default setting of Mplus. We visually assessed the status of the posterior distributions and assessed convergence based on the potential scale reduction factor (PSRF) for each parameter using a threshold of 1.05 (Gelman & Rubin, 1992), which is also the default setting of Mplus; PSRF should be close to 1 if the Markov chains have converged to the target posterior distribution (Gelman & Rubin, 1992). We evaluated model fit based on the Bayesian variant of the root mean square error of approximation (BRMSEA), the Bayesian comparative fit index (BCFI), and the Bayesian Tucker-Lewis index (BTLI) (Hoofs et al., 2018, Garnier-Villarreal & Jorgensen, 2019, Asparouhov & Muthén, 2019). For BCFI and BTLI, Asparouhov and Muthén (2019) suggested criteria that required values above 0.95 for adequate fit. For BRMSEA, Asparouhov and Muthén (2019) proposed three possible outcomes: the fit index is inconclusive (lower limit ≤ 0.06 ≤ upper limit), the index suggests that the model fits reasonably well (both lower and upper limits < 0.06), and the index suggests a poor fit between model and data (both lower and upper limits > 0.06).

After fitting the full model, we trimmed nonsignificant paths from the model until only significant paths remained (Kline et al., 2011). We explored the robustness of the main analysis results by re-fitting the final model for each response variable using datasets of observed distributions rather than the fundamental distributions estimated by Maxent, null model 2, and the two minimum numbers of butterfly species noted above (i.e., 15 and 20) within each grid cell.

## Results

Throughout the Japanese archipelago, butterfly species richness increased toward lower latitudes (south) and higher elevations (Figs. 1a, 2a), and both the mean (Figs. 1b, 2b) and divergence (Figs. 1c, 2c) of butterfly body size increased toward lower latitudes. The Pareto alpha parameter, which indicates the proportion of species with narrower host breadths, increased toward higher latitudes (north) (Figs. 1d, 2d), while the mean (Figs. 1e, 2e) and divergence (Figs. 1f, 2f) of phylogenetic host breadth decreased toward higher latitudes (Table S2).

**Figure 1.**
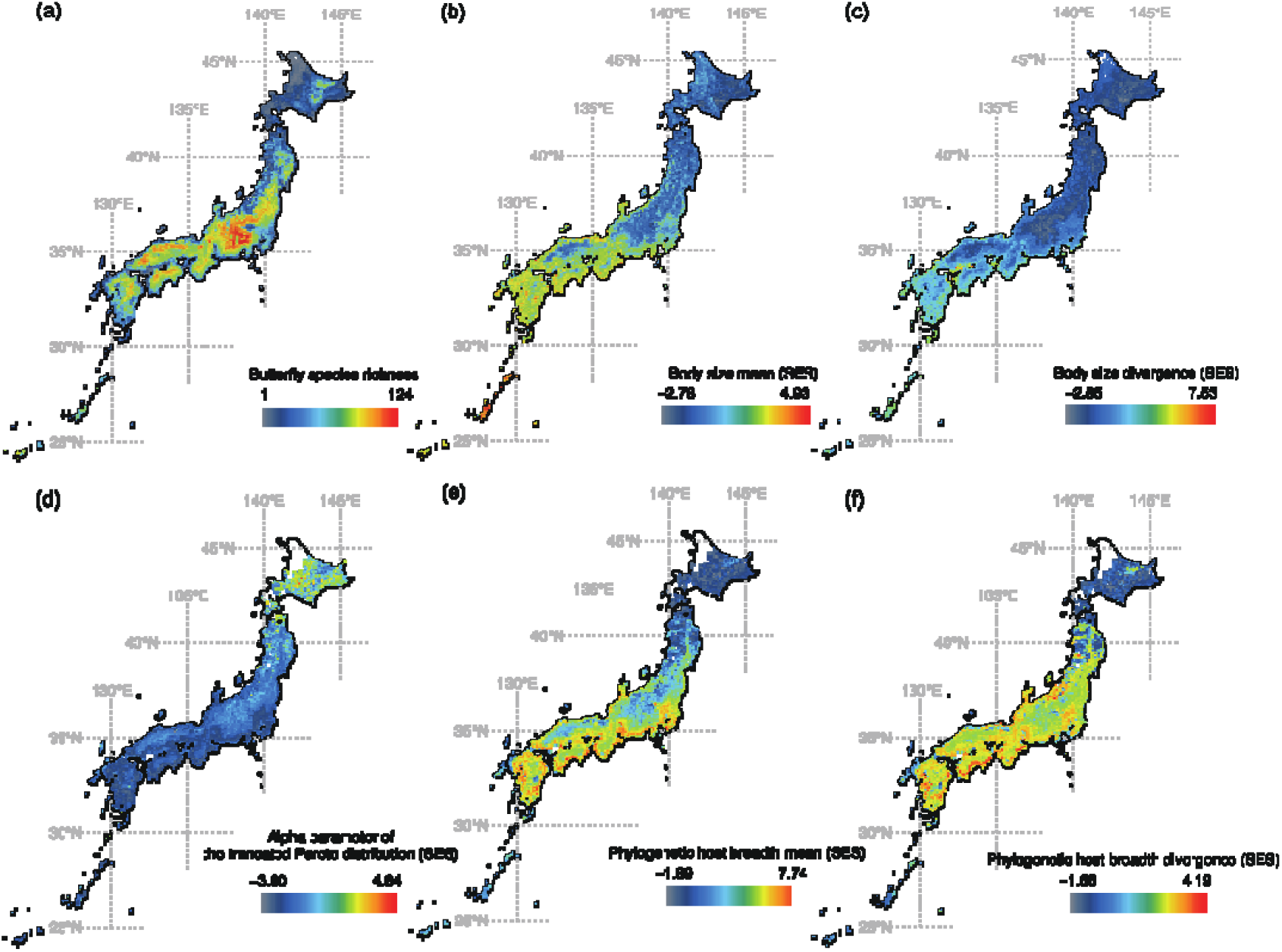
Patterns of (a) butterfly species richness based on stacked predicted species distribution data (Nakadai et al., 2018), (b) mean body size of the butterflies, (c) body size divergence, (d) alpha parameter of the discrete truncated Pareto distribution, (e) mean phylogenetic host breadth, and (f) phylogenetic host breadth divergence across the Japanese archipelago. Only grids that satisfied the criteria (see text) are shown in (d), (e), and (f). Values in all panels except (a) represent standardized effect size (SES) estimates based on null model 1. A dichromatic version of this figure is shown in Figure S5.

**Figure 2.**
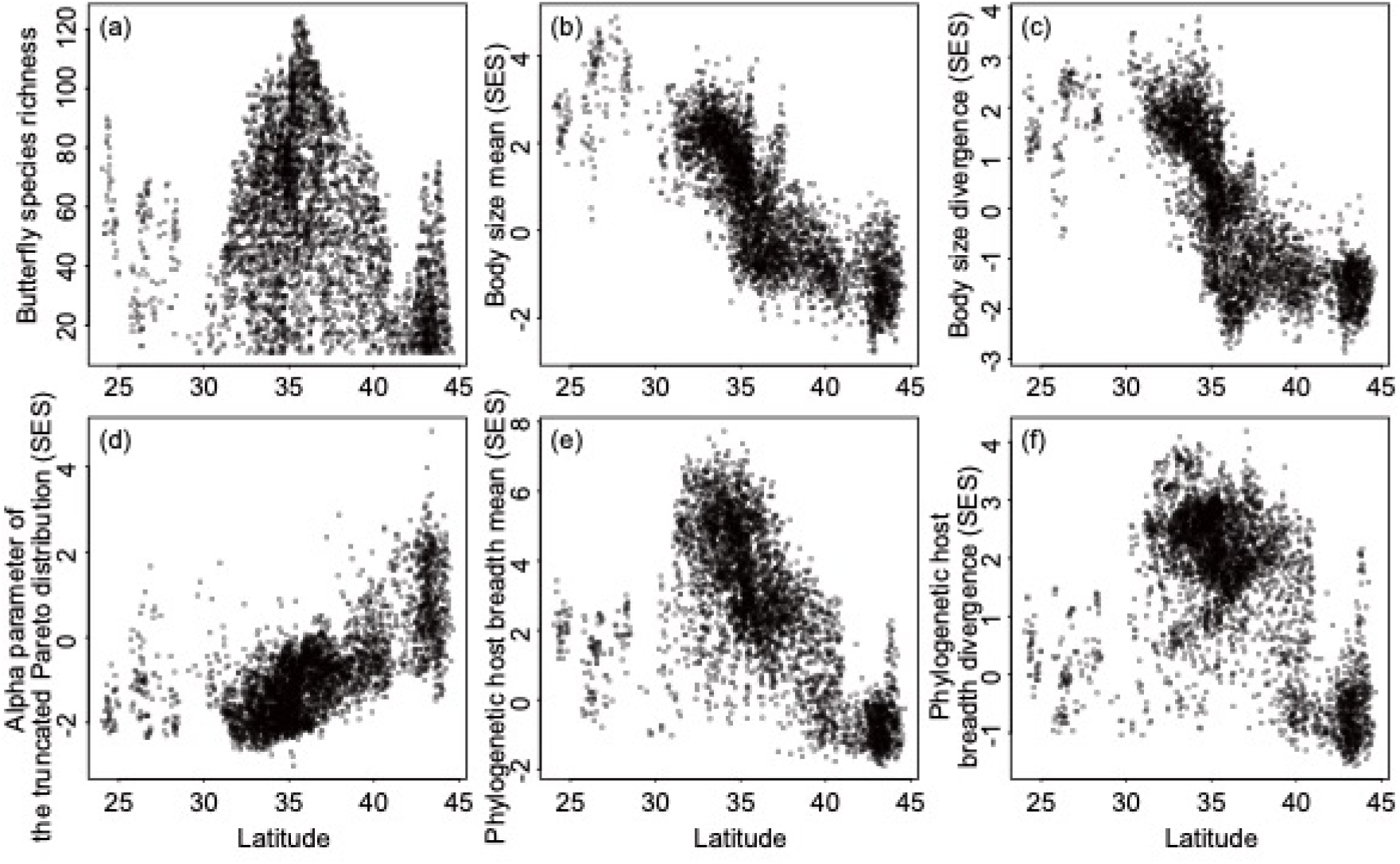
Pairwise associations between latitude and (a) butterfly species richness, (b) mean body size of the butterflies, (c) body size divergence, (d) alpha parameter of the discrete truncated Pareto distribution, (e) mean phylogenetic host breadth, and (f) phylogenetic host breadth divergence across the Japanese archipelago. Values in all panels except (a) represent standardized effect size (SES) estimates based on null model 1. Note that the order of panels in this figure corresponds to that of Figure 1.

Of the 4,668 total grid cells covering the Japanese archipelago, 3909 passed our filtering criteria and were included in the Bayesian SEM (Table 2). The final model satisfied the criteria of good fit, as measured using BCFI, BTLI and BRMSEA (Table 3). Only two of the 52 paths in the full model were nonsignificant, and therefore were removed during the process of model simplification (Table 2).

**Table 2.**
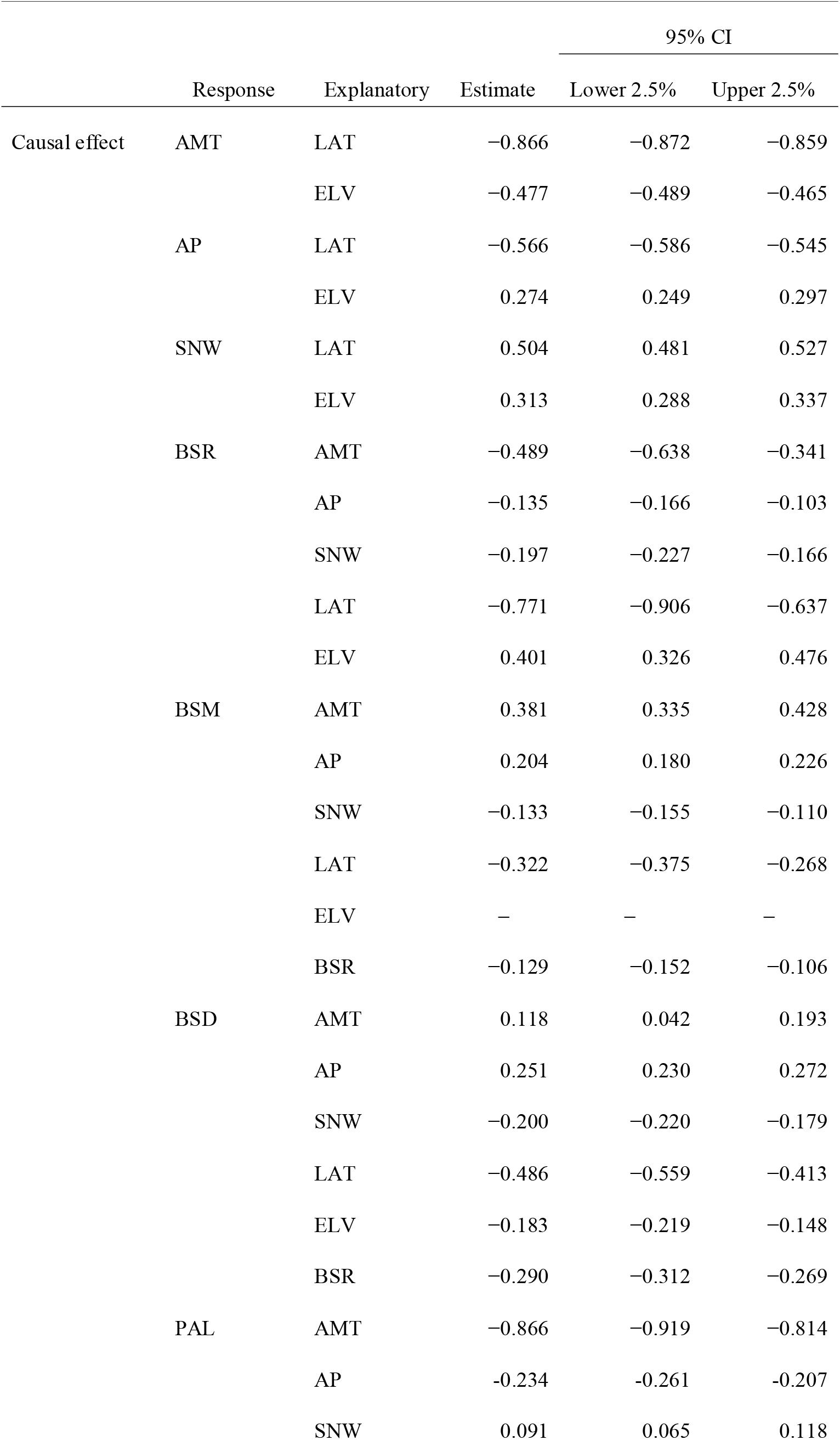
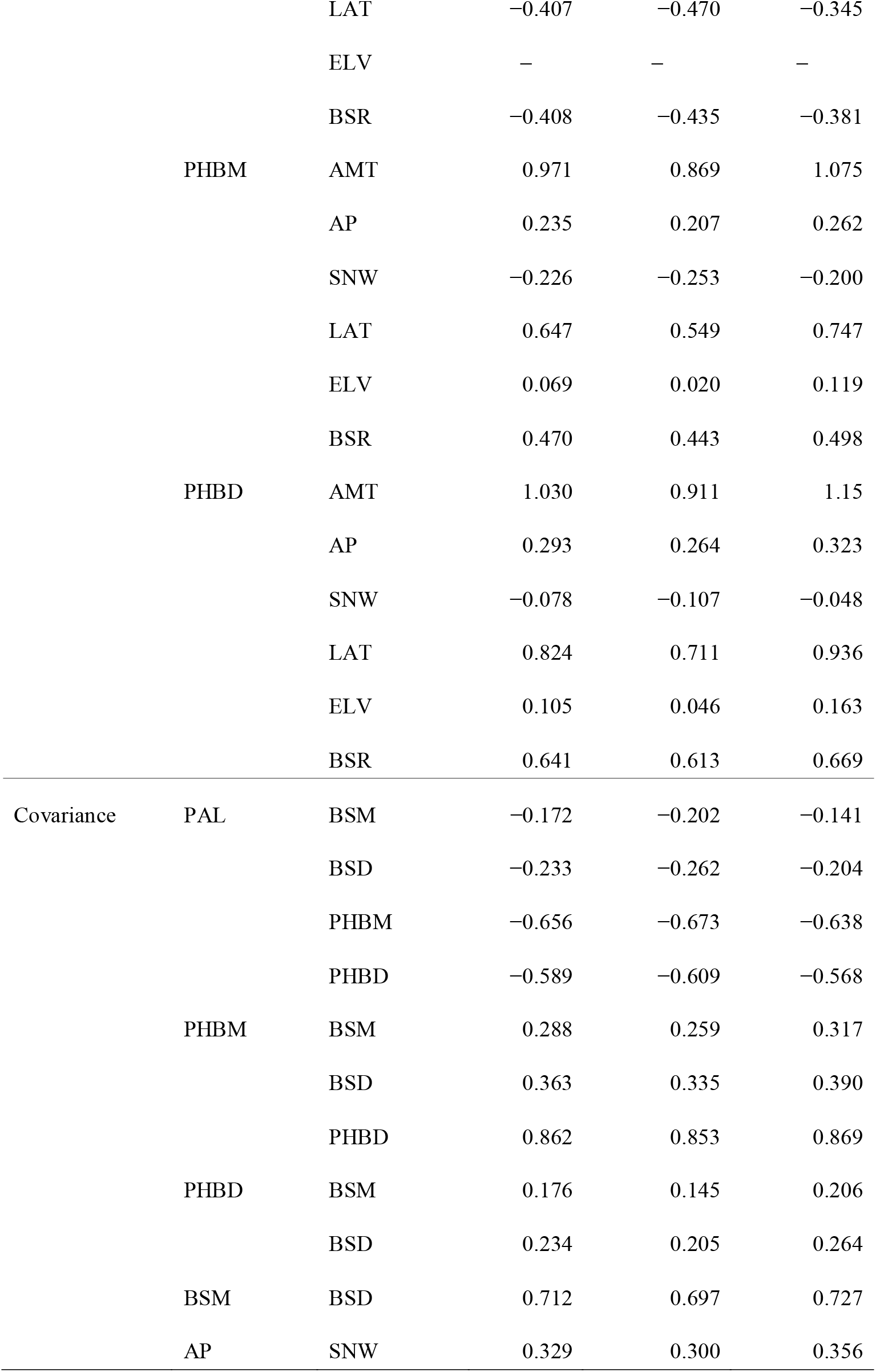
Results of the final Bayesian SEM focusing on the alpha parameter of the discrete truncated Pareto distribution fitted to butterfly host breadth. “Response” and “Explanatory” columns indicate response and explanatory variables, respectively. Parameter estimates with upper and lower limits of their 95% confidence intervals are shown in the next columns (estimates are omitted (–) for statistically non-significant parameters for which the 95% CI overlaps with zero). Abbreviations: LAT: latitude, ELV: elevation, AP: annual precipitation, AMT: annual mean temperature, SNW: annual maximum depth of snow cover, BSR: butterfly species richness, BSM: body size mean, BSD: body size divergence, PAL: alpha parameter of the discrete truncated Pareto distribution, PHBM: phylogenetic host breadth mean, and PHBD: phylogenetic host breadth divergence.

**Table 3.**
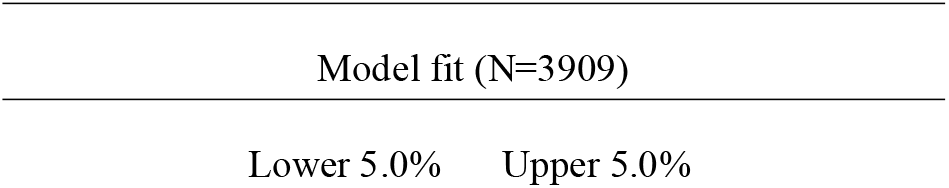
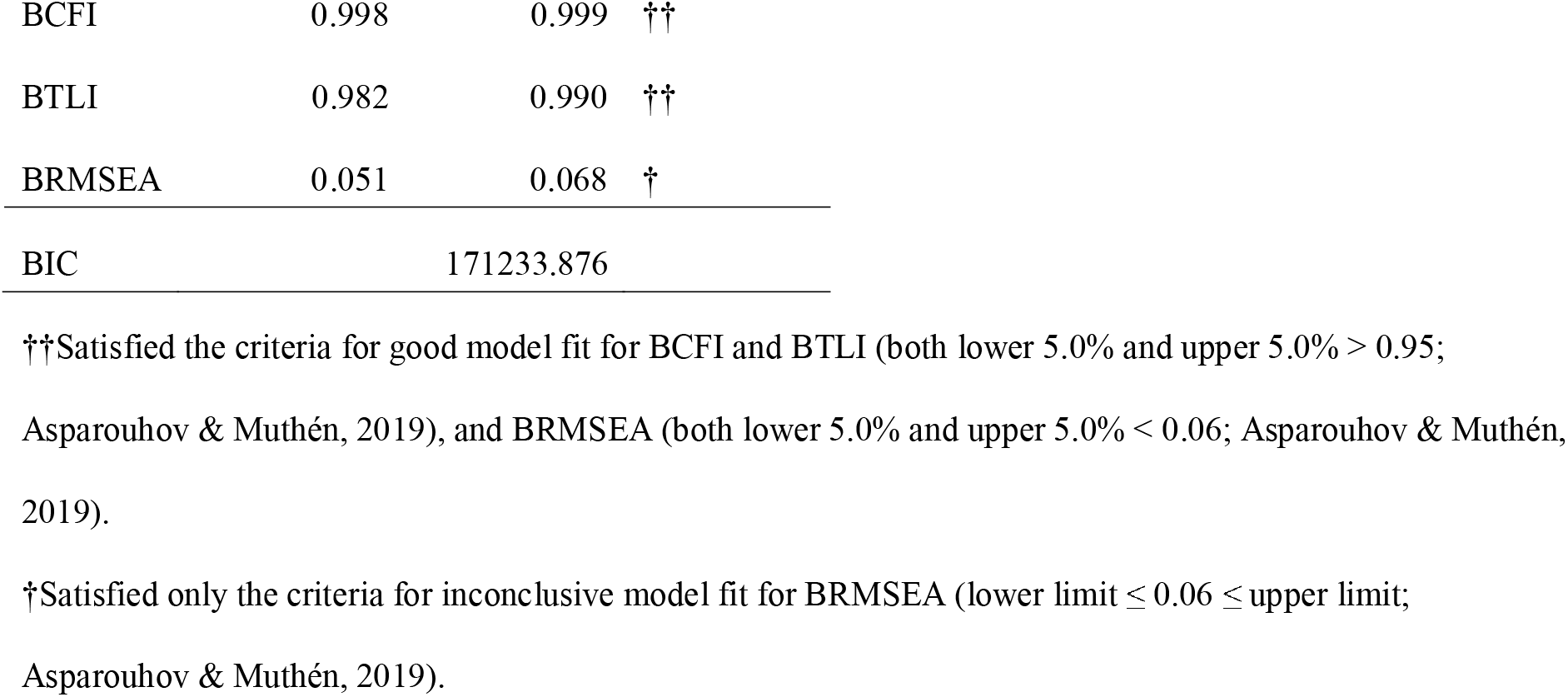
Estimates of the fit of our Bayesian SEM according to different criteria.

| Model fit (N=3909) |  |
| --- | --- |
| Lower 5.0% | Upper 5.0% |

|  |  |  |  |
| --- | --- | --- | --- |
| BCFI | 0.998 | 0.999 | †† |
| BTLI | 0.982 | 0.990 | †† |
| BRMSEA | 0.051 | 0.068 | † |
| BIC | 171233.876 |  |  |

††Satisfied the criteria for good model fit for BCFI and BTLI (both lower 5.0% and upper 5.0% > 0.95; Asparouhov & Muthén, 2019), and BRMSEA (both lower 5.0% and upper 5.0% < 0.06; Asparouhov & Muthén, 2019).
†Satisfied only the criteria for inconclusive model fit for BRMSEA (lower limit ≤ 0.06 ≤ upper limit; Asparouhov & Muthén, 2019).

Of the many factors affecting host breadth, annual mean temperature was most strongly associated with host breadth in the SEM (Table 2, Fig. 3). Specifically, communities containing low concentrations of species with narrow host ranges (i.e., those with low Pareto alpha values) increased with annual mean temperature, as did the mean and divergence for phylogenetic host breadth at the community level (Table 2, Fig. 3). Negative correlations were found between Pareto alpha and the mean and divergence of body size (Table 2, Fig. 3). In addition, positive correlations were identified between the mean and divergence of phylogenetic host breadth and the mean and divergence of body size (Table 2, Fig. 3). Conversely, no significant correlation was found between body size and host breadth at the species level (Fig. S2).

**Figure 3.**
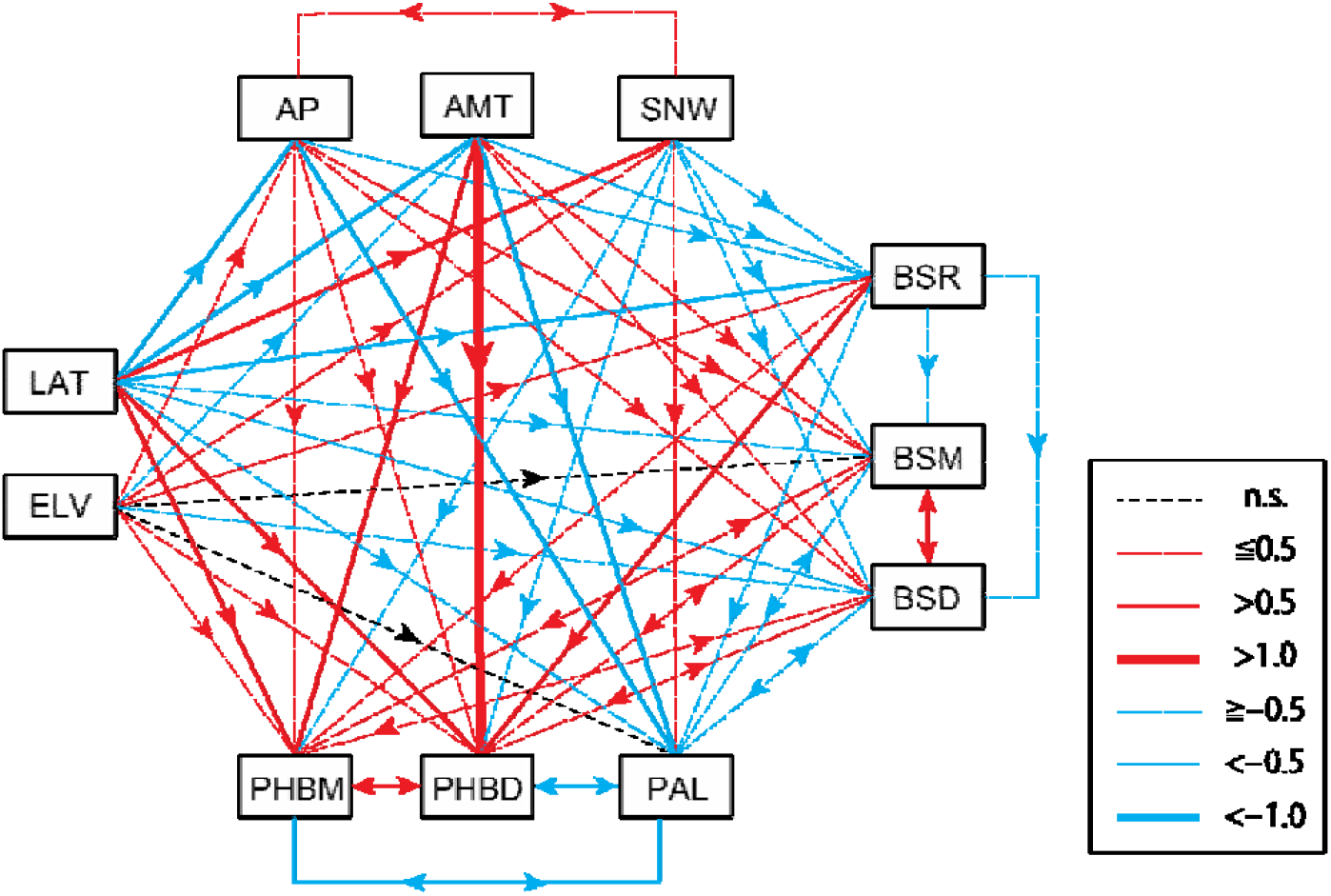
Path diagram of Bayesian SEM explaining host-breadth parameters. Parameter estimates are shown in Tables 2 and S3. Red arrows indicate significant positive effects or correlations, and blue arrows indicate significant negative ones. Line weights indicate the strength of the standardized estimates (see legend). Black dashed lines indicate the paths removed during the modification process. Abbreviations: LAT: latitude, ELV: elevation, AP: annual precipitation, AMT: annual mean temperature, SNW: annual maximum depth of snow cover, BSR: butterfly species richness, BSM: body size mean, BSD: body size divergence, PAL: alpha parameter of the discrete truncated Pareto distribution, PHBM: phylogenetic host breadth mean, and PHBD: phylogenetic host breadth divergence.

Butterfly species richness was driven by annual mean temperature, annual precipitation, annual maximum depth of snow cover, latitude, and elevation. Based on standardized estimates, latitude was the strongest (negative) driver of butterfly species richness, while annual mean temperature was the second most effective (negative) predictor (Table 2, Fig. 3).

The mean of butterfly body size was strongly driven by annual mean temperature (positively) and latitude (negatively), while the effects of annual precipitation, annual maximum depth of snow cover and butterfly species richness were relatively weak (Table 2, Fig. 3). Butterfly body size divergence was driven by latitude, annual mean temperature, annual precipitation, annual maximum depth of snow cover, elevation and butterfly species richness. Of these factors, latitude strongly negatively influenced body size divergence (Table 2, Fig. 3). Furthermore, a positive correlation was found between the mean and divergence of butterfly body size (Table 2, Fig. 3).

The significance and the strength of the major relationships of both host breadth and butterfly body size with the environmental predictors were consistent when the final models were re-run with datasets of the observed distributions (Table S4) or null model 2 (Table S5), and with two alternative criteria for the minimum number of butterfly species per grid cell (Table S6).

## Discussion

### The latitudinal pattern of fundamental host breadth is different than predicted patterns of realized host breadth

One of the most widespread assumptions in ecology is that the degree of resource specialization increases toward lower latitudes (i.e., toward the tropics) (Moles & Ollerton, 2016; Dyer & Forister, 2018). By contrast, our results show that the host breadth of butterfly communities becomes more specialized at higher latitudes (i.e., toward the north) across the Japanese archipelago. Thus, our finding apparently contradicts the widespread assumption of how species combine to form local communities. According to our results, the host breadth of butterfly communities was most strongly associated with climatic factors (i.e., annual mean temperature, annual precipitation, and annual maximum depth of snow cover), a geographic factor (latitude), and butterfly species richness. Specifically, both mean and divergence of host breadth of butterfly communities were higher where temperatures were higher. Furthermore, grid cells with high butterfly diversity showed broader and more divergent host ranges.

The trend of broader and more divergent host ranges within communities at lower latitudes may be produced due to relaxing of physiological constraints with increased energy availability, thereby reducing the relative importance of abiotic environmental filters and allowing for wider variation in niches (Park, 1949; Fischer, 1960). Read et al. (2018) suggested that relaxing of physiological constraints combined with stronger biotic interactions among species toward the tropics leads to increased resource specialization, which is the opposite trend to our findings. As mentioned in the Introduction, this study focused on fundamental host breadth, which was defined as the host breadth when not limited by interspecific interactions within a trophic level. Evidently, without considering interspecific interactions, the relaxation of abiotic regulation toward lower latitudes allows for the occurrence of butterfly species with broader host ranges. Thus, our results appear contrary to the conventional assumption of latitudinal gradients in resource specialization. However, previous studies have failed to clarify the constraints of interspecific interactions on resource specialization among herbivorous insects. In fact, the existence of strong interspecific resource competition among herbivorous insects, which would cause niche partitioning, has repeatedly been questioned (Lawton & Strong, 1981; Strong et al., 1984; Condon et al., 2014; Nakadai & Kawakita, 2017). Furthermore, Nakadai et al. (2018) even showed that butterfly species pairs that use similar host-plant taxa tend to have overlapping distributions in the Japanese archipelago. Thus, the relevance of interspecific competition to the latitudinal gradient in the diet breadth of herbivorous insects is not clear. Also, the southern part of the Japanese archipelago, is more fragmented into small islands than are the northern and middle parts, and resource specialization in the southern part showed lower values than at the southern edge of the Kyusyu island and Honshu. While island size might influence resource specialization via resource availability and effects on body size (reviewed in Palkovancs 2003), assessing such effects was not possible with the present data.

Along the focal latitudinal and climatic gradients, many variables other than host breadth may change together or independently, including both the body size and diversity of butterflies. Teasing apart the directionality of causalities among these patterns is a long-standing research subject in ecology (Wasserman & Mitter, 1978; Loder et al., 1998). Consistent with previous studies on plant-feeding insects (Brown & Maurer 1989; Pöyry et al., 2017), we identified a strong positive relationship between the mean body size and mean host breadth at the grid community level. Lindström et al. (1994) argued that larger body size in lepidopterans allows for a larger “home range,” and thereby increases encounters with a wide range of host plants, but also requires more energy, which is more easily acquired if the species is a broad generalist. In our dataset, both the mean and divergence of body size increased toward lower latitudes and higher temperatures (Table 2). This pattern of diminishing body sizes toward higher latitudes is referred to as a converse Bergmann cline (although most interspecific studies on insects have found no clines; Shelomi, 2012). The southward increases in both the mean and divergence of body size are consistent with the hypothesis based on relaxed physiological constraints (Table 1). Furthermore, mean body size is positively correlated with mean phylogenetic host breadth. Increasing mean body size within a community may therefore be one of the factors enhancing diet generalization in communities of herbivorous insects (Table 1). At higher latitudes or elevations, shorter growing seasons also limit the duration of the larval stage, restricting the distributions of large-bodied or slow-developing insects (Blanckenhorn & Demont, 2004; Shelomi, 2012). For example, Zeuss et al. (2017) reported that, among European lepidopteran species, average body size and the average number of generations per year decrease with latitude. However, we did not detect significant correlations between body size and host breadth at the species level (Fig. S2). Therefore, the relationship between mean body size and mean phylogenetic host breadth at the grid level may simply reflect correlations driven by the responses of both variables to the same underlying environmental predictors (e.g., annual mean temperature).

### Potential drivers of contrasting patterns between fundamental and realized resource specialization

The largest difference between our analytical approach and those of previous studies concerns the properties in the herbivore community dataset. The analyses of Forister et al. (2015) and many others (e.g., Novotny et al., 2006) were based on direct observations of insect–plant associations in local communities. By contrast, our ‘species-pool’ approach based on modeled distributions may combine the distributions of some species pairs that would typically not occur at the same site at the same time, for reasons such as phenological niche division. Our analyses did not focus on phenological patterns along the latitudinal gradient, as our main focus was on “fundamental” patterns at a large spatial scale. If the difference between our results and those of other studies (e.g., Forister et al., 2015) are due to differences in the approach between fundamental and realized resource specialization, biotic filtering across trophic levels (i.e., tri-trophic interactions; reviewed in Schemske et al., 2009) may be an important local-scale factor (Cavender-Bares et al., 2009). For example, narrow and overlapping host breadths may be strongly promoted by host-specific natural enemies (e.g., Condon et al., 2014), and enemy pressure is thought to be greater in the tropics (Roslin et al., 2017; Romero et al., 2018). Local adaptation of herbivorous butterflies is a frequent phenomenon (e.g., Singer & Parmesan, 1993; reviewed in Gandon & Van Zandt, 1998), and locally specialized enemies associated with host plants and variation in plant resistance to herbivory (i.e., coevolutionary interactions) may result in large variations in host plants among local sites, in turn leading to wider host breadths in focal species at the regional level. It seems possible for a species to have high local specialization in a lower latitude community, even if its members tend to have a relatively broad host breadths throughout their range. Over time, this should translate into diversification among these specialized local populations, creating the expected positive latitudinal gradient in host breadth. However, because the Japanese archipelago does not include stable tropical regions, it is unclear whether there has been enough time for local plant species to adapt to the full butterfly fauna, or a subset thereof. In the Japanese archipelago, geographic patterns of realized resource specialization of herbivorous butterflies have never been investigated, so we unfortunately could not evaluate the existence of differences between fundamental and realized resource specialization. However, in future works, comparisons between fundamental and realized resource specialization through standardized sampling methods in the same systems would be highly beneficial for understanding the specific mechanisms that drive differences in specialization across taxa and regions.

### Methodological considerations

To elucidate the detailed drivers of host breadth in Japanese butterflies, we attempted to include as many factors as possible in our analyses. However, our results also reveal several important limitations of current methods that must be addressed in future studies. Unfortunately, we could not directly evaluate the effects of regional plant diversity on the geographic distribution of resource specialization. Plant diversity is highest near the center (at circa N 35°) of the Japanese archipelago (e.g., Kubota et al., 2015). In fact, Kubota et al. (2015) demonstrated that overall vascular plant richness and endemism are highest in central Japan, and that elevation and annual mean temperature are major drivers of the pattern, indicating that the trend is driven by more than just latitude. Central Japan is characterized by mountainous regions, and elevation may be an indirect predictor of plant diversity. In fact, the patterns of the simple correlations in our study indicated that elevation is strongly correlated with butterfly species richness but is less correlated with host breadth (Table S2). Therefore, plant diversity in itself does not appear to be a strong driver of fundamental host breadth.

In general, the proportion of rare plant species increases toward the tropics (e.g., Ulrich et al., 2016), possibly due to negative density-dependent survival effects (Janzen–Connell hypothesis; Janzen, 1970). Such processes were suggested as a fundamental explanation for the contrasting latitudinal pattern (i.e., more generalists toward lower latitudes) of resource specialization found in the simulation results of Forister and Jenkins (2017). Specifically, a high proportion of rare plants would favor the evolution of greater host breadth in insects, causing the mean and divergence of host breadth to increase toward lower latitudes (Ghazoul & Sheil, 2010). The evenness of the biomass of plant species is a potential driver of resource specialization, although the evenness of plant abundance (e.g., number of individuals) was used as an explanatory variable in previous simulation studies (Forister & Jenkins 2017). In any case, data on both plant biomass and abundance were not available for use in our study or in previous studies. Therefore, investigations focusing on the available biomass of resources are needed to further explore the effects of plant community composition on herbivore host breadth. Furthermore, we were unable to include phylogenetic information on butterfly species in our analysis. Both life history traits and distributions may be affected by evolutionary history (i.e., phylogenetic conservatism). This is a limitation of both our study and previous investigations and may have introduced bias. Phylogenetic information hopefully can be incorporated into models in the future, when the lepidopteran tree of life becomes better resolved.

In addition, we focused mainly on the transitional region between the hemiboreal and subtropical zones, and the patterns observed may not be generalizable to the global scale. Regional studies focusing on narrower latitudinal gradients should consider geohistorical effects (e.g., when the region was colonized by particular species or where hotspots of endemism are located). Our findings also may differ from the patterns observed in previous studies (e.g., Forister et al., 2015) because of our more limited geographic scope, i.e., the absence of large tropical regions, which have significant in-situ diversification rates. The effects of co-evolutionary interactions may be absent as there has been relatively little in-situ diversification within our study area. In fact, most butterfly species on the Japanese archipelago are not endemics but rather are widely distributed across the Eurasian continent. This pattern is largely explained by the fact that the Japanese archipelago was repeatedly connected to the continent with land bridges throughout glacial stages of the Pliocene and Pleistocene epochs, allowing for butterfly dispersal (details reviewed in Tojo et al., 2017). On the other hand, detailed regional studies have the advantage of providing extensive and accurate information on host use. Previous studies have suggested that sampling effort directly affects the inferred degree of host specialization (Dixon et al., 1987), but such effects are difficult, if not impossible, to address in a large-scale meta-analysis.

### Historical and evolutionary perspectives

We did not address evolutionary processes in this study, as phylogenetic information was not available for all of the studied butterfly species. However, our findings should be generalizable to other regions and broader spatial patterns, as they are based on evolutionary processes. For example, we could expand on the evolutionary concept of species association dynamics recently proposed by Nylin et al. (2017) and the oscillation hypothesis proposed by Janz et al. (2001, 2006), which focus on speciation and species distributions, by integrating spatial perspectives by adding a latitudinal direction of oscillation dynamics. The oscillation hypothesis proposed by Janz et al. (2001, 2006) postulates that, over time, herbivorous insect lineages oscillate between a generalist state, during which they may expand their geographical ranges, and various specialist states, which present outcomes of local adaptation of individual populations. In this model, speciation occurs after host specialization. To explore this issue further, we plotted minimum and maximum values of phylogenetic host breadth and body size within butterfly communities against latitude (Fig. S3), and confirmed that the ranges of these trait values become increasingly narrow toward higher latitudes. If physical constraints on host breadth or body size in insect herbivores exist (Fig. S3), very large or very small butterfly species should have difficulty extending their distribution ranges from the south to the north, and therefore such extreme species would be filtered out. Our findings are consistent with the proposal of Fischer (1960), who argued that the stable climate in the tropics allows for a wider range of morphological and physiological adaptations than can occur in temperate regions. On the other hand, the spread of butterfly species with broader host ranges from south to north might be hindered by biogeographic limitations (Kimura, 2004; Hirao et al., 2015). Therefore, we propose a novel prediction: butterfly species with large host breadths are more likely to have expanded their distributions to the south or to have originated in the south and then specialized and diversified there via oscillation processes, and only descendant species with narrower host ranges could migrate northward. Although biogeographic limitations and abiotic factors (e.g., temperature, precipitation, and snow) are important determinants of butterfly distributions (Quinn et al., 1998; Luoto et al., 2006), our predictions provide further insights into the importance of geographical and morphological constraints to species association dynamics and the oscillation hypothesis. These predictions may also be combined with general ecological principles such as Rapoport’s rule (Rapoport, 1982), which is related to latitudinal distribution range and global climate history and their dramatic effects on species distributions (Nyman et al., 2012). Specifically, Rapoport’s rule would alter the distributional range of each species, which may affect the possibility of local resource specialization via northward migration. In addition, climatically driven distributional changes along a north– south axis, as well as associated changes in the abundance of different plant groups, have been proposed as factors influencing the patterns and processes of herbivore diversification (Nyman et al., 2012). Therefore, the inclusion of dynamic distributional changes along a latitudinal gradient may generalize the oscillation hypothesis for understanding the mechanisms that drive biodiversity.

## Conclusions

Butterfly communities become more specialized in their host use toward higher latitudes (i.e., toward the north) across the Japanese archipelago, a pattern that is opposite of the predictions arising from classic hypotheses. Our analyses clearly revealed how the variation in resource specialization is distributed, as well as the potential influence of physiological constraints resulting from variation in body size along the same geographical gradient. Since the work of Ehrlich and Raven (1964), coevolutionary interactions between plants and herbivores have been considered the main determinant of host breadth. Furthermore, our results highlight the importance of the current climate as a factor regulating butterfly host breadth and morphology (Fischer, 1960), regardless of whether the impact is direct or indirect. Future studies should explore how latitudinal gradients in the phylodiversity and evenness of herbivores and plant resources influence community-level host breadth. We emphasize that the approach used here is applicable to many other areas and taxa for which reliable information on species occurrences and niches are available. The continuous improvement of public databases on species distributions and host use (e.g., Global Biodiversity Information Facility and HOSTs: Robinson et al. 2010) will hopefully eventually allow testing of patterns of fundamental resource specialization on a global scale. Importantly, comparative analyses of fundamental and realized resource specialization in communities can allow important insights into factors that determine the assembly and local realizations of eco-evolutionary interaction networks across the globe.

## Data accessibility statement

Information on the forewing lengths of Japanese butterflies was published by Nakadai et al. (2020). Environmental and geographic data were obtained from Mesh Climate Data 2010 (Japan Meteorological Agency, http://nlftp.mlit.go.jp/ksj/gml/datalist/KsjTmplt-G02.html) and the National Land Numerical Information database (Geospatial Information Authority of Japan), respectively. The host plant dataset was taken from Saito et al. (2016). Occurrence data for butterfly species are available from the website of the Biodiversity Center of Japan, Ministry of the Environment (http://www.biodic.go.jp/index_e.html).

## Acknowledgements

We are particularly grateful to the local entomologists and naturalists who collected the butterfly data that we compiled for use in this study. We also thank Dr. Matthew Forister, who provided R code associated with the alpha shape parameter. The Biodiversity Center of Japan and the Japanese Ministry of the Environment kindly provided us with datasets on butterfly distributions. Dr. Yasuhiro Sato provided insightful comments on an early version of the manuscript. Financial support was provided by the Japanese Society for the Promotion of Science (grant numbers 15J00601 and 18J00093, to R.N.) and the Academy of Finland (grant number 324392, to A.V.).

## Biosketch

Ryosuke Nakadai is a research associate in National Institute for Environmental Studies, with broad interests in biodiversity and biological interactions across temporal and spatial scales. His works include analysis of biodiversity patterns and development of both statistical and molecular tools for a wide range of taxa.

## Author contributions

RN conceived of and designed the study and analyzed the data; RN and TN constructed the figures; KH, RN, and TI compiled the butterfly dataset; RN and AV wrote the first draft; and all authors significantly contributed to the final text.

## Appendices

Figure S1 Geographical patterns of climatic variables and elevation: (a) annual mean temperature, (b) annual precipitation, (c) annual maximum depth of snow cover, and (d) elevation. A dichromatic version of this figure is shown in Figure S6.

Figure S2 Relationships of species-specific body size with (a) the number of host-plant genera (*r* = −0.062, *p* = 0.156) and (b) phylogenetic host breadth (*r* = −44.82, *p* = 0.131) across Japanese butterflies. Points indicate species. Significant correlations were not confirmed by linear regressions.

Figure S3 Latitudinal patterns of (a) minimum phylogenetic host breadth (*r* = –0.022, *p* < 0.001), (b) maximum phylogenetic host breadth (*r* = –0.072, *p* < 0.001), (c) minimum body size (*r* = 0.020, *p* < 0.001), and (d) maximum body size (*r* = –0.167, *p* < 0.001). Values represent standardized effect sizes (SES) estimated using null model 1. A dichromatic version of this figure is shown in Figure S7.

Figure S4 Latitudinal patterns of (a) the alpha parameter of the discrete truncated Pareto distribution based on predicted occurrence data (*r* = 0.018, *p* < 0.001), and (b) the alpha parameter of the discrete truncated Pareto distribution based on observed occurrence data (*r* = 0.005, *p* < 0.001). Values represent non-standardized trends comparable to those of Forister et al. (2015).

Figure S5 Patterns of (a) butterfly species richness based on stacked predicted species distribution data (Nakadai et al., 2018), (b) mean body size of the butterflies, (c) body size divergence, (d) alpha parameter of the discrete truncated Pareto distribution, (e) mean phylogenetic host breadth, and (f) phylogenetic host breadth divergence across the Japanese archipelago. Only grids that satisfied the criteria (see text) are shown in (d), (e), and (f). Values in all panels except (a) represent standardized effect size (SES) estimates based on null model 1. The full color version is shown in Figure 1 of the main text.

Figure S6 Geographical patterns of climatic variables and elevation: (a) annual mean temperature, (b) annual precipitation, (c) annual maximum depth of snow cover, and (d) elevation. The full color version is shown in Figure S1 of the main text.

Figure S7 Latitudinal patterns of (a) minimum phylogenetic host breadth (*r* = –0.022, *p* < 0.001), (b) maximum phylogenetic host breadth (*r* = –0.072, *p* < 0.001), (c) minimum body size (*r* = 0.020, *p* < 0.001), and (d) maximum body size (*r* = –0.167, *p* < 0.001). Values represent standardized effect sizes (SES) estimated using null model 1. The full color version is shown in Figure S3 of the main text.

Table S1 Species analyzed in the study. Nine species were excluded from the analyses for various reasons. First, the taxonomic status of three species changed during the interval between the fourth and fifth biodiversity censuses (Inomata, 1990; Shirôzu, 2006); we excluded those species because the identifications were unreliable. Second, we excluded three non-herbivorous species (including an omnivore, *Shirozua jonasi*) and three gymnosperm-feeding species.

Table S2 Correlations between environmental variables and calculated indices.

Table S3 Comparison of the results between the full model and final model.

Table S4 Comparison of the results based on predicted and observed distribution datasets. The model structures are described in Table 2.

Table S5 Comparison of the results based on null model 1 and null model 2. The model structures are described in Table 2.

Table S6 Comparison of the results based on different criteria of minimum butterfly species richness. The model structures are described in Table 2.

Table S7 Environmental variables and calculated indices that were used in this study for all 10 km grids.

