## Supplementary figures and images for "Fundamental resource specialization of herbivorous butterflies decreases toward lower latitudes"

### Figs_S1-7.pdf

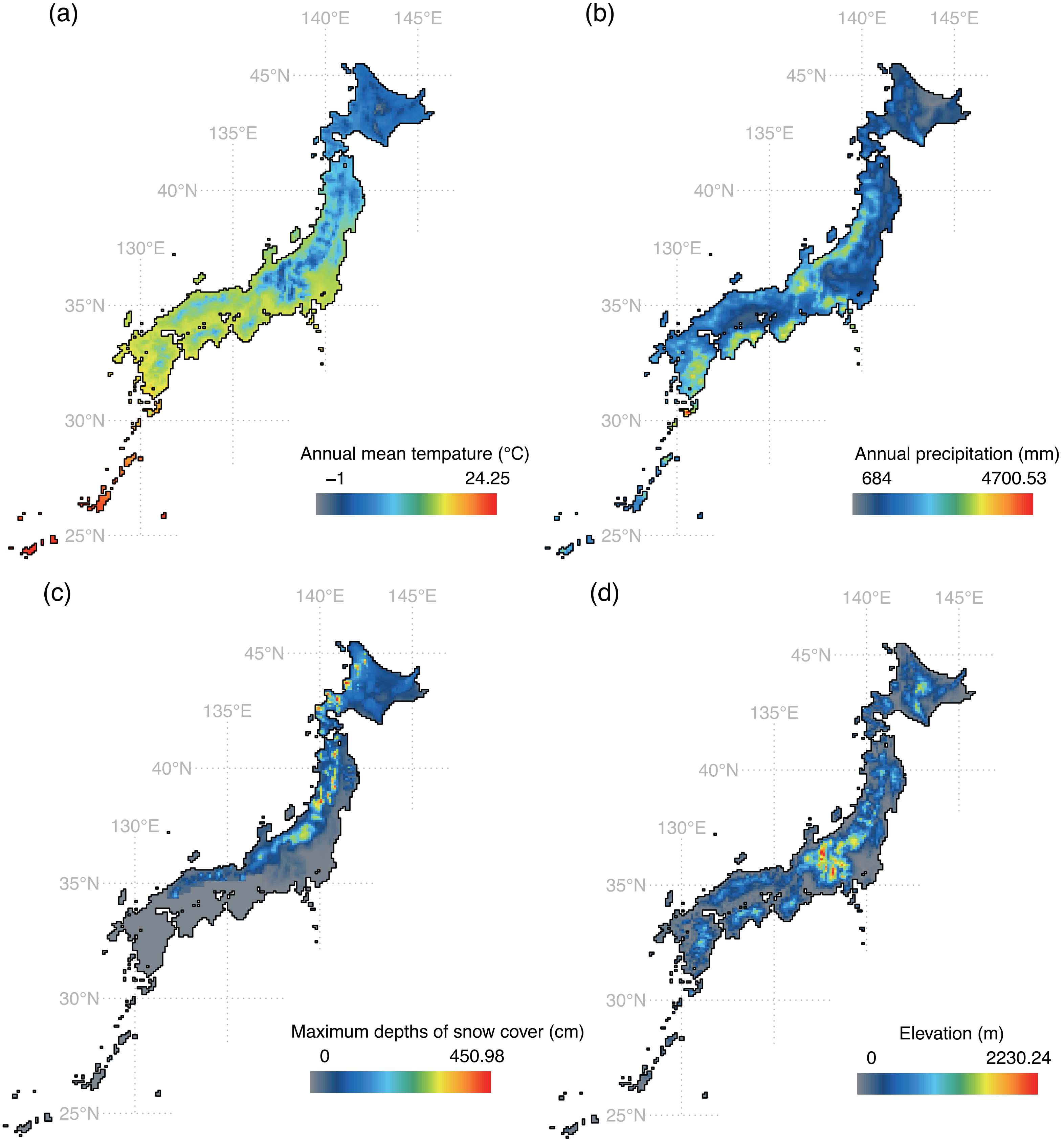

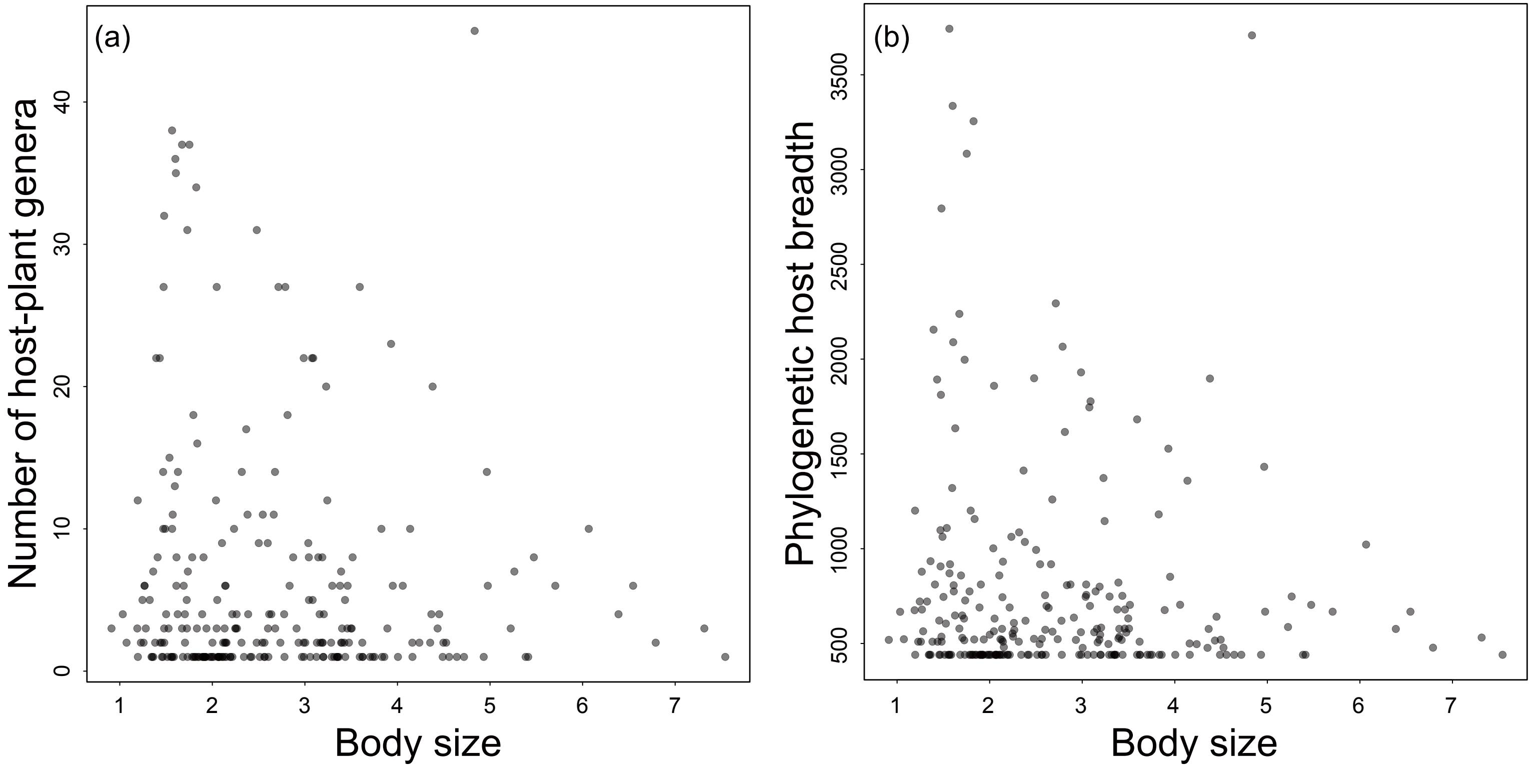

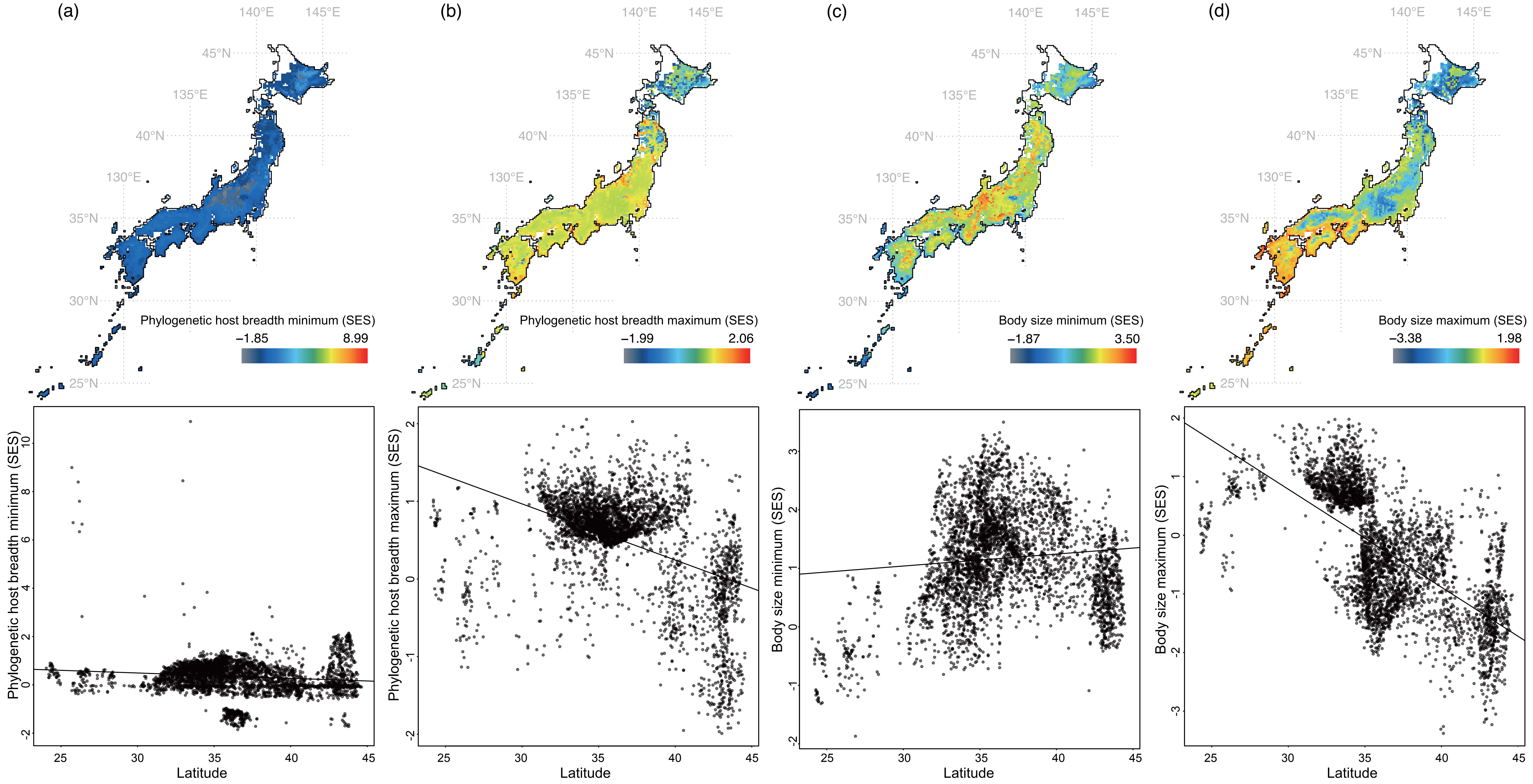

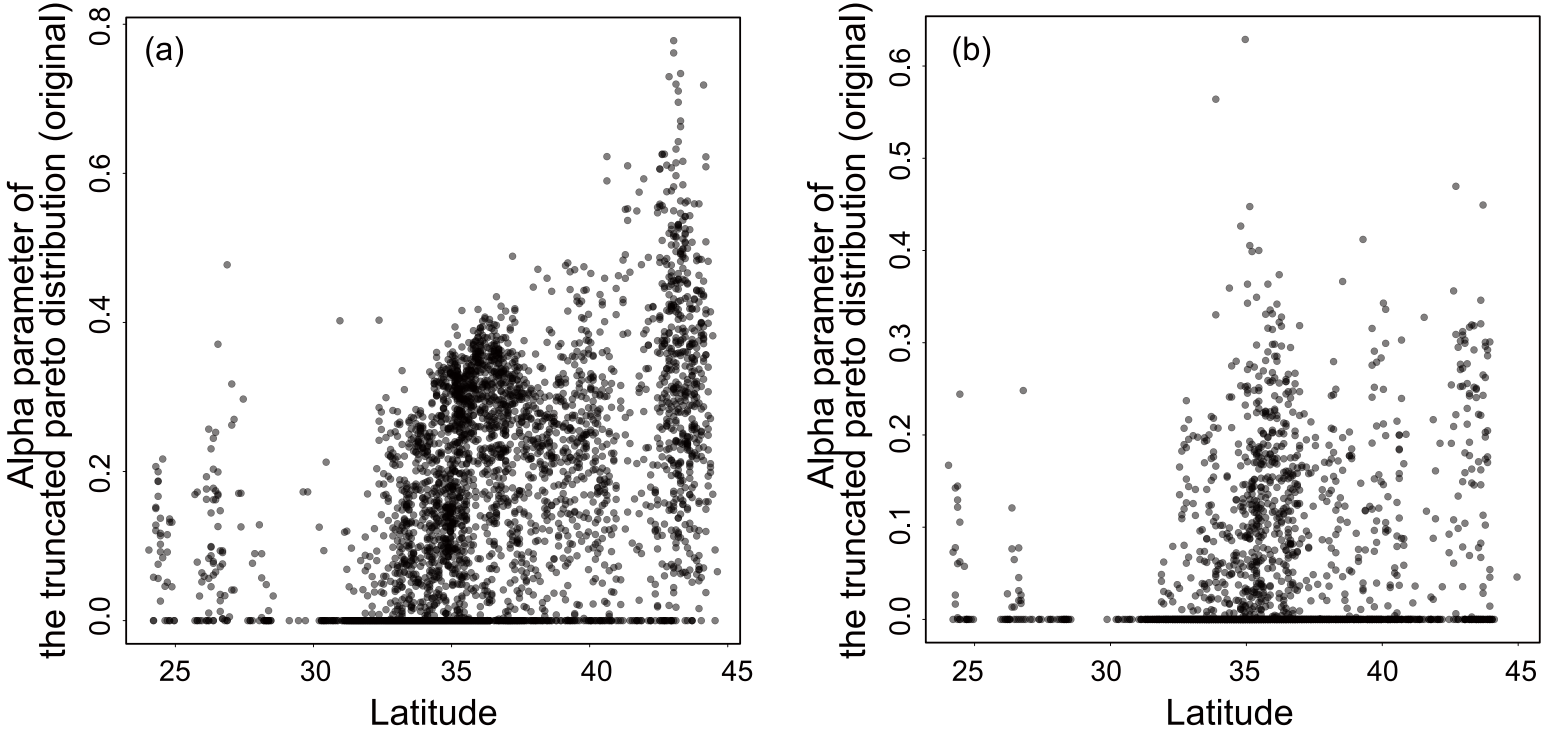

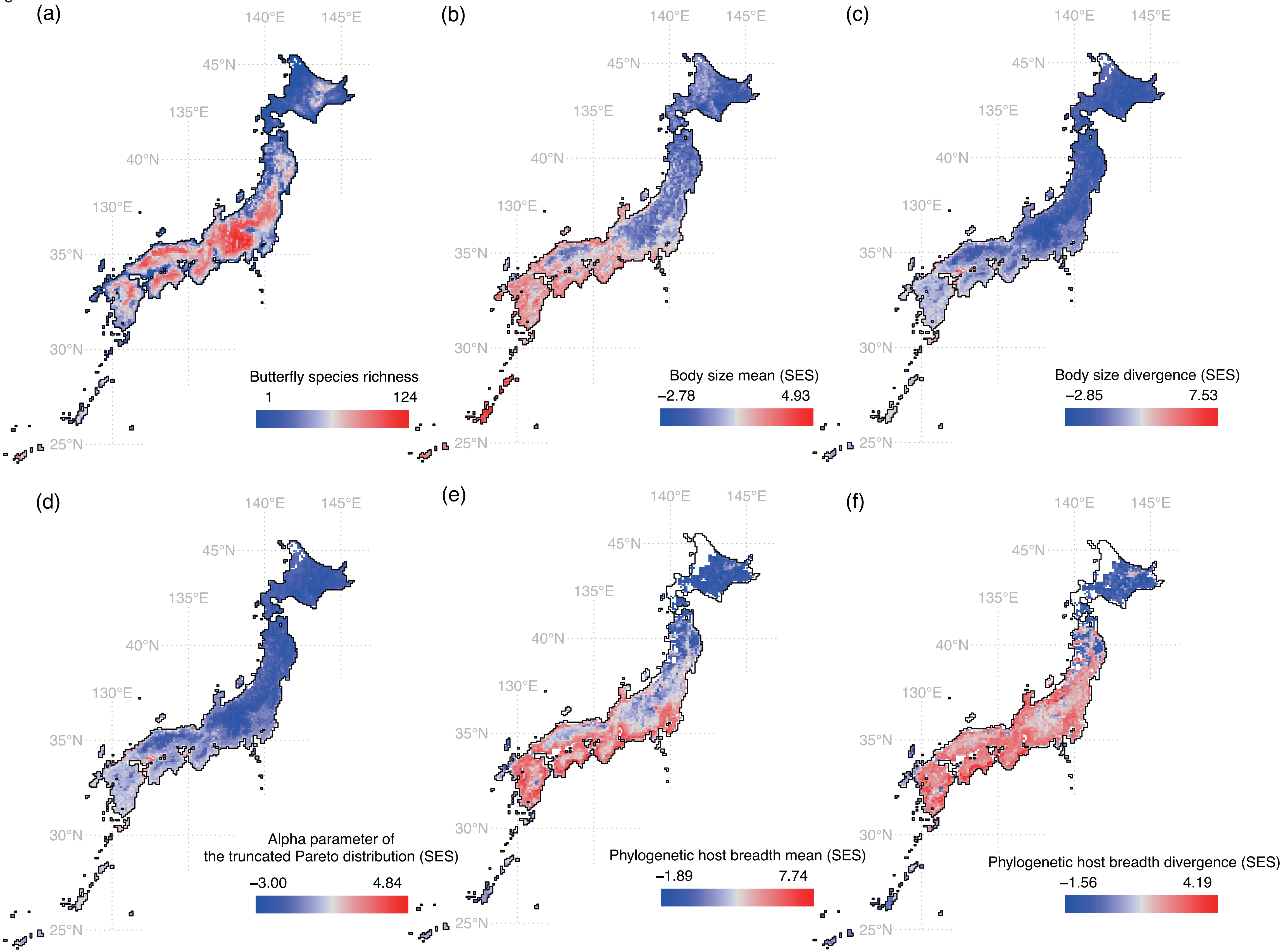

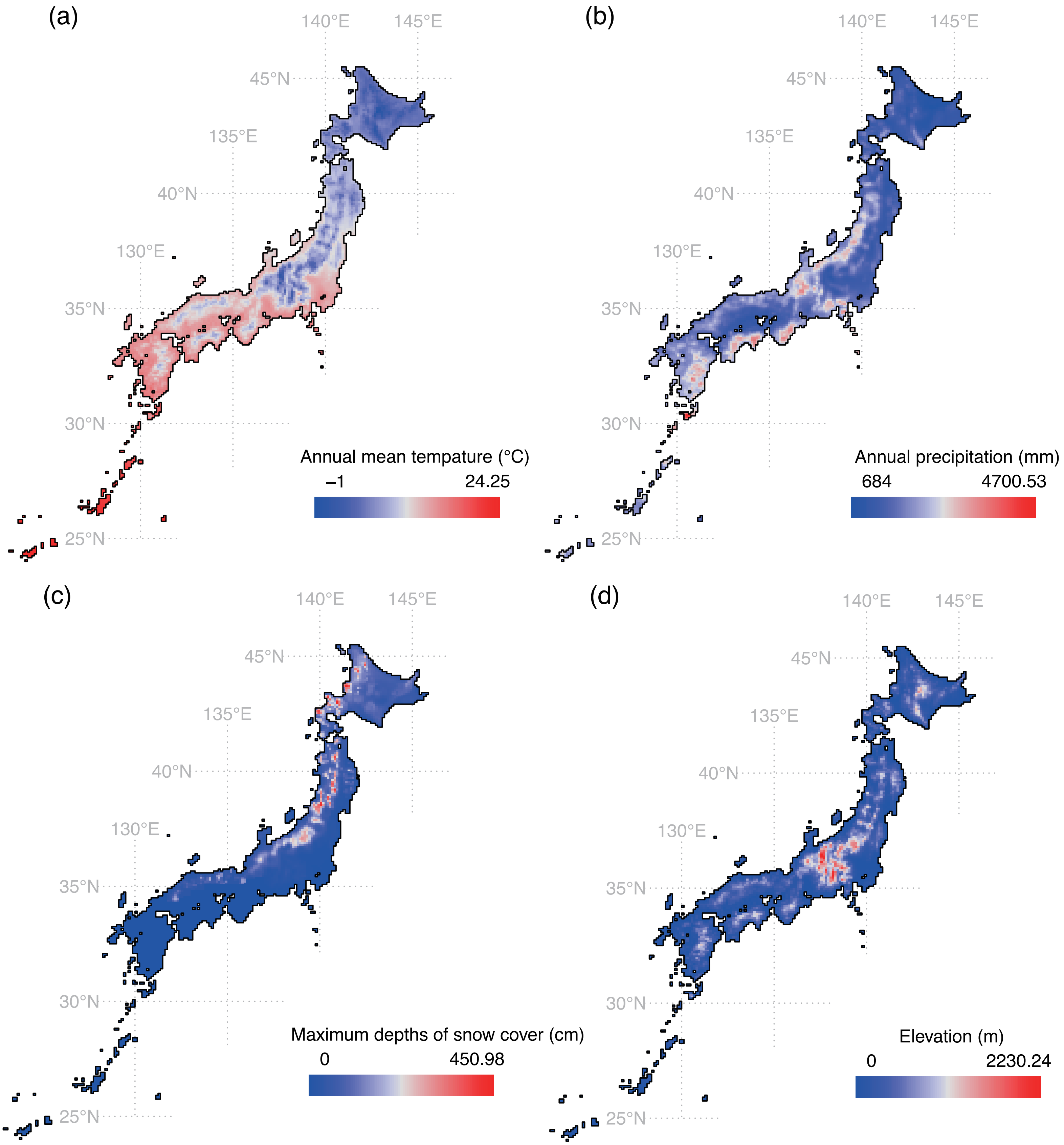

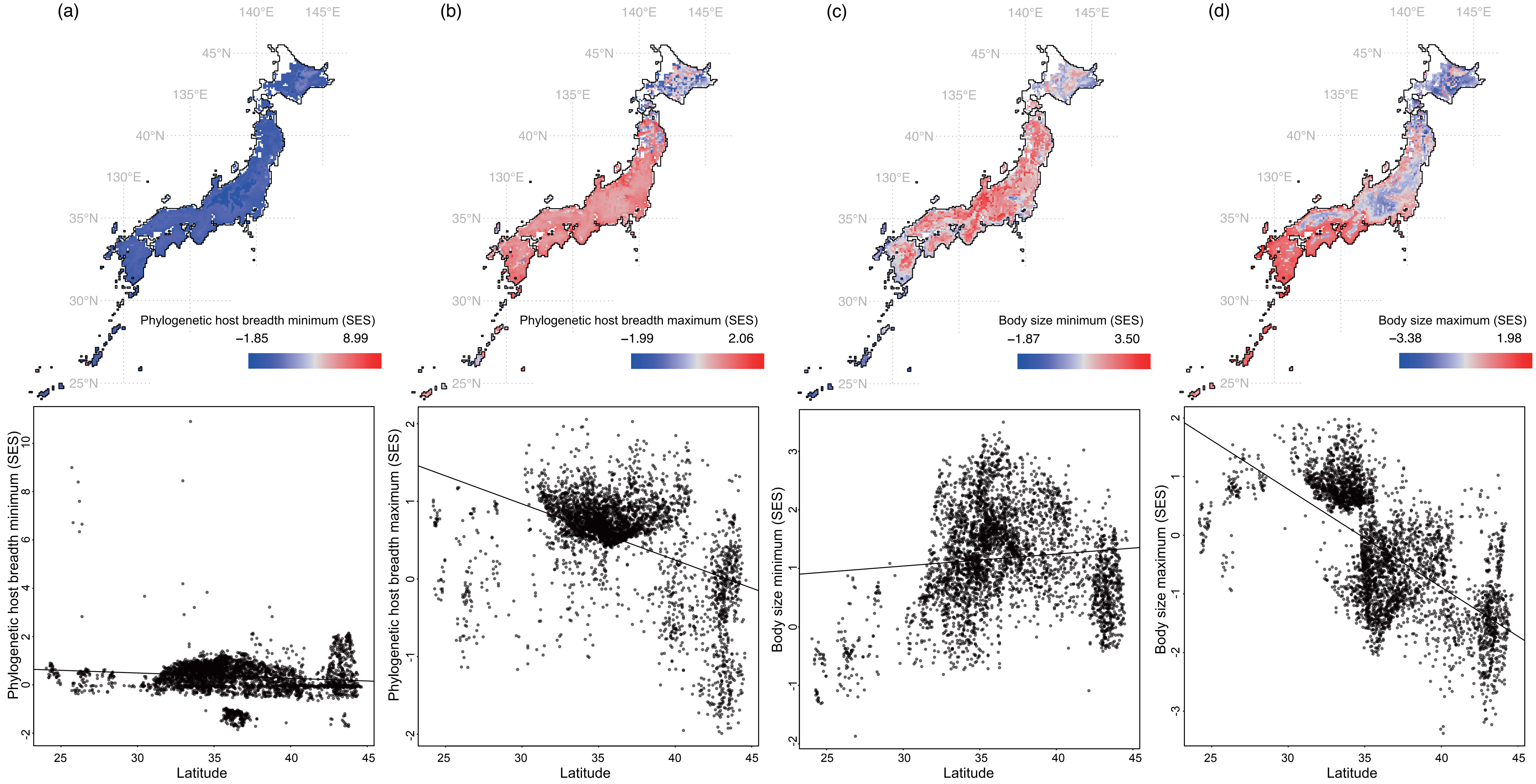
